# The functional convergence and heterogeneity of social, episodic, and self-referential thought in the default mode network

**DOI:** 10.1101/753509

**Authors:** Tanya Wen, Daniel J Mitchell, John Duncan

## Abstract

The default mode network (DMN) is engaged in a variety of cognitive settings, including social, semantic, temporal, spatial, and self-related tasks. Andrews-Hanna et al. (2010, 2012) proposed that the DMN consists of three distinct functional-anatomical subsystems – a dorsal medial prefrontal cortex (dMPFC) subsystem that supports social processing and introspection about mental states; a medial temporal lobe (MTL) subsystem that contributes to memory retrieval and construction of mental scenes; and a set of midline core hubs that are involved in processing self-referential information. We examined activity in the DMN subsystems during six different tasks: (1) theory of mind and (2) moral dilemmas (for social cognition), (3) autobiographical memory and (4) spatial navigation (for memory-based construction/simulation), (5) self/other adjective judgement (for self-related cognition), and finally, (6) a rest condition compared to a working memory task. At a broad level, we observed similar whole-brain activity maps for the six contrasts, and some response to every contrast in each of the three subsystems. In more detail, both univariate analysis and multivariate activity patterns showed partial functional separation, much of it in close accord with the proposals of separate dMPFC and MTL subsystems, though with less support for common activity across anterior and posterior regions of a midline core. Integrating social, spatial, self-related, and other aspects of a cognitive situation or episode, multiple components of the DMN may work closely together to provide the broad context for current mental activity.

**Significance Statement:** Activity in the default mode network (DMN) can been found across a wide range of high-level tasks that involve social, semantic, episodic, or self-referential cognition. Given this diversity, an important proposal is that the DMN can be parcellated into subsystems with different cognitive functions. The current experiment employed a wide range of experimental tasks to directly test for functional convergence and heterogeneity between DMN regions. The results support both partial differentiation and integration; working together, distributed DMN regions may assemble the multiple contextual components of a cognitive situation or episode.

## Introduction

The default mode network (DMN) was originally discovered as a collection of medial prefrontal, lateral temporal, lateral parietal, and posterior medial cortical regions that reliably exhibit enhanced activity during passive rest compared to simple, externally oriented tasks (Shulman et al., 1997; Raichle et al., 2001). Raichle et al. (2001) postulated that the DMN is involved in cognitive states that are suspended during many attentionally-demanding tasks. A large body of literature has now provided evidence that the DMN supports several aspects of spontaneous and deliberate self-generated thought that transcend the immediate sensory environment (Christoff et al., 2004, 2009; Buckner et al., 2008; Andrews-Hanna, 2012; Andrews-Hanna et al., 2014b). Complementing this strong activity during rest, subsequent work has shown DMN activity across a variety of high-level tasks, including social (Greene and Haidt, 2002; Mars et al., 2012; Molenberghs et al., 2016), semantic (Binder et al., 2009; Humphreys and Lambon Ralph, 2017), episodic (Ranganath and Ritchey, 2012; Rugg and Vilberg, 2013), and self-referential (Kelley et al., 2002) cognition.

One common proposal is that the DMN represents broad features of a cognitive episode, scene or context (Hassabis and Maguire, 2007; Ranganath and Ritchey, 2012; Manning et al., 2014; Crittenden et al., 2015; Baldassano et al., 2017; Smith et al., 2018). This episode might be imagined, as in spontaneous mind-wandering or recollection of a previous event, or currently perceived (Ranganath and Ritchey, 2012; Manning et al., 2014; Baldassano et al., 2017; Smith et al., 2018). Contextual representations might include spatial, social, temporal, self-related and other features, with reduced processing of these features during focused attention on the details of an external task, but enhancement during spontaneous, self-generated cognition at rest.

A core question is the degree of heterogeneity across DMN regions. Early reviews (Buckner and Carroll, 2007; Buckner et al., 2008), meta-analyses (Spreng et al., 2009), as well as experimental data (Spreng and Grady, 2010) suggested that spatial, social, memory and imagination tasks produce substantially overlapping DMN activity. More recently, consistent with the multiple features of a cognitive context, some studies suggest that the DMN exhibits heterogeneous functional components (Andrews-Hanna et al., 2010, 2014a; Andrews-Hanna, 2012). In an important synthsis, Andrew-Hanna et al. (2010) partitioned the DMN into three subsystems. A dorsal medial prefrontal cortex (dMPFC) subsystem, composed of the dorsal medial prefrontal cortex (dMPFC), the temporoparietal junction (TJP), the lateral temporal cortex (LTC), and the temporal pole (TempP), is involved in “introspection about mental states”, including theory of mind, moral decision making, social reasoning, story comprehension, and conceptual processing. A medial temporal lobe (MTL) subsystem, consisting of the ventromedial prefrontal cortex (vMPFC), the posterior inferior parietal lobe (pIPL), the retrosplenial cortex (RSC), the parahippocampal cortex (PHC), and the hippocampal formation (HF+). subserves “memory-based construction/simulation”, including autobiographical memory, episodic future thinking, contextual retrieval, imagery, and navigation. These two subsystems are proposed to converge on a midline core, consisting of the anterior prefrontal cortex (aMPFC) and the posterior cingulate cortex (PCC). The core subserves valuation of “personally significant information”, as well as linking social and mnemonic processes shared with the dMPFC and MTL subsystems.

The current study further investigates separation and integration across the DMN. To this end we examined patterns of univariate and multivoxel activity across six tasks, aiming to separate social cognition, memory-based construction/simulation, self-related cognition and rest. Across this combination of tasks and analysis methods, we found a degree of functional separation between DMN regions, largely consistent with the Andrews-Hanna (2010) dMPFC and MTL subsystems, though less so with their concept of the midline core. To a degree, however, we also found overlapping activity across the whole DMN, with each task producing some activation in each subsystem. While subsystems of the DMN system appear somewhat specialized, our data also suggest collaboration in assembling the multiple components of a cognitive situation or context.

## Methods

### Participants

27 participants (13 male, 14 female; ages 20-39, mean = 24.8, SD = 4.3) were included in the experiment at the MRC Cognition and Brain Sciences Unit. An additional participant was excluded due to excessive head motion (> 5 mm). All participants were fluent English speakers, neurologically healthy, right-handed, with normal or corrected-to-normal vision. Participants were also required to be familiar with navigating in Cambridge city centre. Procedures were carried out in accordance with ethical approval obtained from the Cambridge Psychology Research Ethics Committee, and participants provided written, informed consent before the start of the experiment.

### Stimuli and task procedures

This study consisted of six tasks that were previously found to engage the DMN. These tasks were: a theory of mind task, a moral dilemmas task, an autobiographical memory task, a spatial imagery task, a self/other adjective judgement task, and a comparison of rest with working memory (Figure 1). For the first five tasks, each run contained two conditions (one condition that has been associated with DMN activity and a matched control condition), along with periods of fixation between trials or blocks. Conditions were presented in randomized order, with the restriction of a maximum of two consecutive trials or blocks of the same condition. For the working memory task, each run contained alternating periods of working memory and periods of fixation. In all runs, participants were instructed to relax and clear their minds of any thought during fixation periods, and fixation periods were jittered and sampled from a random uniform distribution (see details below for each task). Before entering the scanner, participants practiced a shortened version of each task (containing 1∼2 trials or blocks of each condition). Participants were also asked to practice writing down digital numbers until they were able to write all of them in the correct format, and to clarify that they were familiar with all 20 landmark locations used in the spatial imagery task. Inside the scanner, there were two scanning runs for each task. Run order was randomized with the constraint that repeats of the same task were between four and seven runs apart. Before the start of each run, participants were played audio-recorded task instructions to remind them of what to do during that run. Each run lasted approximately 5∼7 minutes.

**Figure 1.**
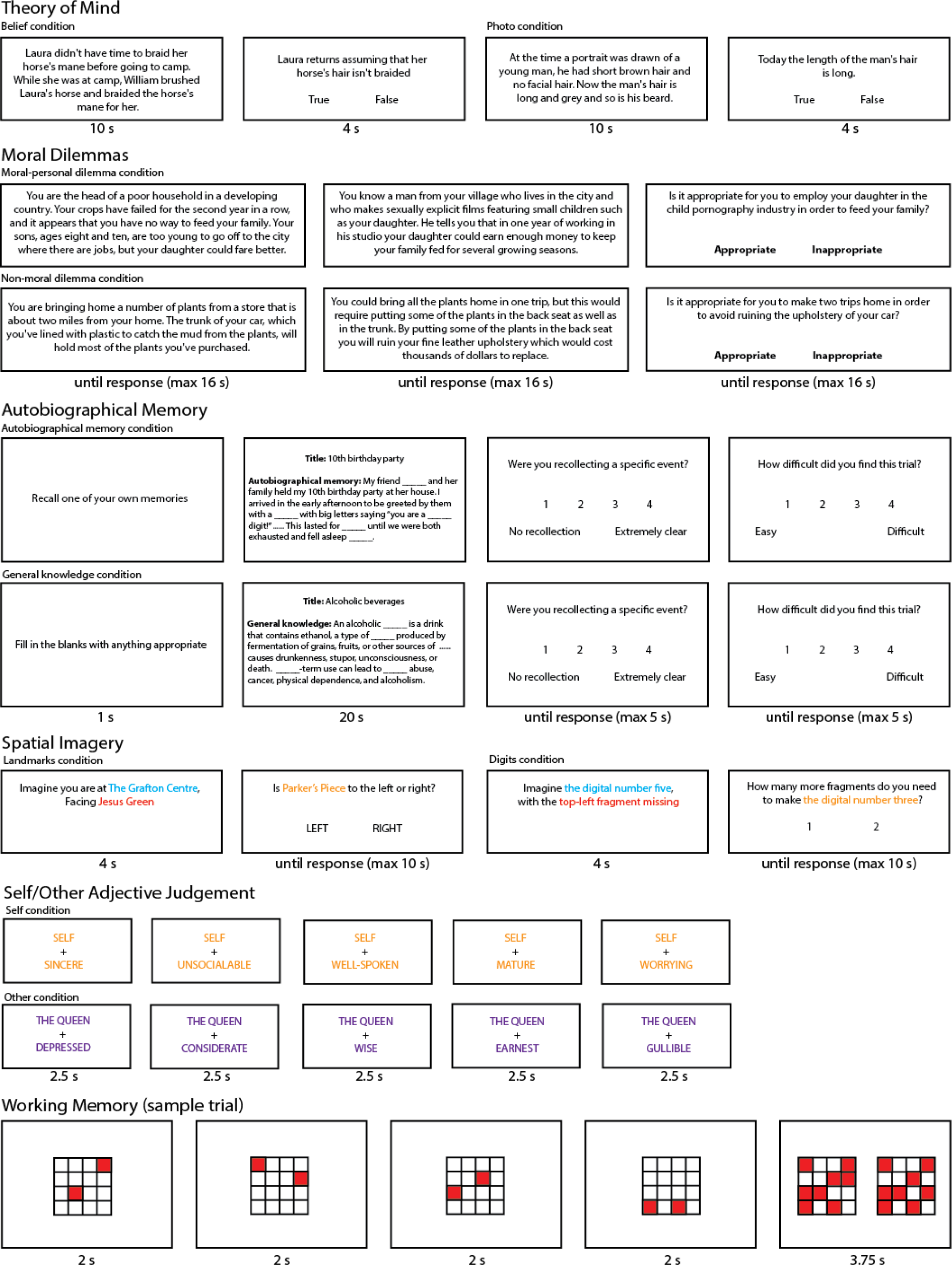
Example trial/block from each of the six tasks: theory of mind, moral dilemmas, autobiographical memory, spatial imagery, self/other adjective judgement, and working memory. All stimuli were shown on a 1920 × 1080 screen (stimulus size and width/height ratio has been adjusted for this figure for illustration purposes). The examples in the autobiographical memory task were shortened to fit in the figure.

All tasks were coded and presented using the Psychophysics Toolbox (Brainard, 1997) in Matlab 2014a (The MathWorks, Inc.). Stimuli were projected on a 1920 × 1080 screen inside the scanner, and participants indicated their responses using a button box, with one finger from each hand in tasks that had two-choice decisions (all tasks except autobiographical memory). According to Andrews-Hanna (2012), the chosen tasks would be hypothesized to differently engage the dMPFC and MTL subsystems, with all tasks engaging the core hubs. The theory of mind and moral dilemmas tasks were chosen as tasks requiring “introspection about mental states” and were hypothesized to recruit the dmPFC subsystem. The autobiographical memory and spatial imagery tasks were chosen as tasks that required “memory-based construction/simulation” and were hypothesized to recruit the MTL subsystem. The self/other judgement task was chosen as a task that involved “personally significant information”, and was hypothesized to recruit predominantly the core hubs. Finally, the working-memory task was chosen to examine the activity of the DMN during passive rest compared to an external task.

#### Theory of Mind task

The theory of mind task was adapted from Dodell-Feder et al. (2010). On each trial, participants were presented with a short story to read for 10 seconds. Afterwards participants were given a statement related to the story and were asked to judge whether it was ‘true’ or ‘false’ by pressing a button (left or right). Some trials involved making judgements about other people’s beliefs, while others involved making non-belief judgements. Each question stayed on the screen up to ten seconds, or until the participant made a button press. This was then followed by a 10∼24 second fixation period before the next trial began. Each run consisted of five trials of each condition (belief and non-belief).

#### Moral Dilemmas task

The moral dilemmas task was adapted from Greene et al. (2001). On every trial, participants were presented with a hypothetical situation that posed a dilemma, which could either be a moral-personal dilemma or a non-moral dilemma. Each dilemma was presented as text through a series of three displays, with the first two describing a scenario and the third posing a question about the appropriateness of an action one might perform in such a situation. The maximum time one display could be on screen was 16 seconds, but when participants finished reading the text, they were allowed to press any button to move on to the next display. On the third display, participants made the appropriateness judgement by pressing a button (left or right). They were told that there was no correct answer for many of the questions, and were asked to consider each situation carefully and provide their best answer. A 6∼8 second fixation cross was presented in between each trial. Each run consisted of five trials of each condition (moral-personal dilemma (MPD) or a non-moral dilemma (NMD)).

#### Autobiographical Memory task

Prior to the experiment, participants were asked to provide twelve written personal memories, each with a title that provided a general description of its contents. Participants were given specific instructions to provide clear memories, where they were able to remember the people, objects, and location details featured in the corresponding memory. Each memory was required to be between 100∼150 words long. All events were required to be temporally and contextually specific, occurring over minutes or hours, but not more than one day. The memories were then edited by the experimenter such that 13∼17 critical words were removed and replaced with a blank underscore line. Occasionally, if the memories participants sent were too long, they were shortened; or if the memories were too short or contained too few details, a new sentence with a prompt was added (e.g., “I was wearing a _____”, “It was around _____o’clock”, “I felt very _____”).

During the task, on a given trial, participants were given a 100∼150-word long text to read with 13∼17 critical words missing, and were asked to fill in the blanks in their mind. Half of the trials used text adapted from the participants’ autobiographical memories; the other half contained text related to general knowledge (either procedural tasks, such as “how to make chocolate chip cookies”, or knowledge about a common topic, such as “alcoholic beverages”). Before the onset of the text display, a 1 second cue was presented to indicate the upcoming condition. For autobiographical memory trials, participants were told to try to “really get into the memory” while filling in the blanks. They were asked to try to imagine themselves reliving that experience. In the general knowledge condition, participants were asked to fill in the blanks with anything appropriate, and to try to “think carefully for good answers”. All trials were terminated after 20 seconds. However, participants were told that there was no need to rush to try to finish all the blanks, and it was more important to be accurate than fast. This was designed to encourage participants to be engaged as much as possible throughout the 20 seconds. After the 20 seconds were over, participants were given two rating questions (‘Were you recollecting a specific event?’ and ‘How difficult did you find this trial?’). They were given 5 seconds to provide each rating on a scale of 1 to 4. Since it involved four buttons, participants gave responses with the four fingers of their right hand. This was then followed by an 8∼12 second fixation period between trials. There were five trials of each condition (autobiographical memory and general knowledge) in each run.

#### Spatial Imagery task

In the spatial imagery task, there were two types of mental imagery conditions, each presented in blocks of trials. One type of block involved judging relative locations of landmarks in Cambridge (this task was adapted from Vass & Epstein, 2017). On each trial, there was first a four second instruction to imagine standing at the landmark indicated in the first line (e.g., Botanic Garden) while facing the landmark indicated in the second line (e.g., King’s College). Afterwards, participants were shown a second screen with a new landmark location (e.g., Parker’s Piece), and were asked to indicate whether it would be on their left or right (in this example the correct answer would be right). The question stayed on the screen for up to 10 seconds, or until participants made a button press. The other type of block involved judging how many fragments were needed to complete a target digital number. At the beginning of each trial, a four second instruction was given to imagine a digital number indicated in the first line (e.g., three) with either an additional fragment or a fragment missing indicated in the second line (e.g., top-right fragment missing). Afterwards, participants were shown a new screen indicating a new target digit (e.g., five), and were asked how many more fragments would need to be added to their original mental image to complete the target (in this example the correct answer would be 1). Participants had up to 10 seconds to answer 1 or 2 (left and right buttons). The two conditions (landmarks and digits) were presented in blocks of four trials, with a 6∼16 second fixation period in between each block. There were four blocks of each condition per run.

#### Self/Other Adjective Judgement task

The self/other judgement task was adapted from Kelley et al. (2002). A total of 160 adjectives were selected from a pool of normalized personality trait adjectives (Anderson, 1968). Half of the words were positive traits and half were negative. On each trial, participants were asked to make a yes/no judgement via button press to indicate whether an adjective shown on the bottom of the screen described the person indicated on the top of the screen (self or the Queen). Each trial was presented for a fixed period of two seconds followed by a 0.5 second fixation. The task was grouped into blocks according to “self” and “the Queen”, with each block consisting of five trials. There were eight blocks of each condition per run. A 6∼16 second fixation period separated each block.

#### Working Memory task

The working memory task was adapted from Fedorenko et al. (2013). On each trial, participants were presented with four consecutive displays. Each display was a 4 × 4 grid, with two of the cells colored red and the remaining white. Each display was presented for two seconds. Afterwards, participants were presented with two choice displays, on the left and right of the screen, one of which had eight red cells in locations corresponding to those from the previous four displays, while the other was similar but with one cell misplaced. Participants were given 3.75 seconds to indicate the correct display by pressing left or right. This was followed by a 0.25 second feedback on the accuracy of their choice. There was a 12∼16 second fixation period between trials. Each run consisted of 16 trials.

### fMRI data acquisition and preprocessing

Scanning took place in a 3T Siemens Prisma scanner with a 32-channel head coil. Functional images were acquired using a standard gradient-echo echo-planar imaging (EPI) pulse sequence (TR = 2000 ms, TE = 30 ms, flip angle = 78°, 64 × 64 matrices, slice thickness = 3 mm, 25% slice gap, voxel size 3 mm × 3 mm × 3 mm, 32 axial slices covering the entire brain). The first five volumes served as dummy scans and were discarded to avoid T1 equilibrium effects. Field maps were collected at the end of the experiment (TR = 400 ms, TE = 5.19 ms / 7.65 ms, flip angle = 60°, 64 × 64 matrices, slice thickness = 3 mm, 25% gap, resolution 3 mm isotropic, 32 axial slices). High-resolution anatomical T1-weighted images were acquired for each participant using a 3D MPRAGE sequence (192 axial slices, TR = 2250 ms, TI = 900 ms, TE = 2.99 ms, flip angle = 9°, field of view = 256 mm × 240 mm × 160 mm, matrix dimensions = 256 × 240 × 160, 1 mm isotropic resolution).

The data were preprocessed and analyzed using automatic analysis (aa) pipelines and modules (Cusack et al. 2014), which called relevant functions from Statistical Parametric Mapping software (SPM 12, http://www.fil.ion.ucl.ac.uk/spm) implemented in Matlab (The MathWorks, Inc., Natick, MA, USA). EPI images were realigned to correct for head motion using rigid-body transformation, unwarped based on the field maps to correct for voxel displacement due to magnetic-field inhomogeneity, and slice-time corrected. The T1 image was coregistered to the mean EPI, and then coregistered and normalized to the MNI template. The normalization parameters of the T1 image were applied to all functional volumes. Spatial smoothing of 10 mm FWHM was applied for whole-brain univariate second-level analysis, but no smoothing was applied for ROI-based analyses or multi-voxel pattern analysis.

A general linear model (GLM) was estimated per participant and per voxel for each of the six tasks. A high-pass filter with 1/128Hz cutoff was applied to both the data and the model. For the first five tasks, regressors were created for each condition, with fixation periods serving as implicit baseline. In the working memory task, one regressor was created for the fixation periods to model passive fixation as the contrast against active task as implicit baseline. Error trials (only applicable for the theory of mind and spatial imagery tasks) and no-response trials were modelled using a separate regressor. All regressors were created by convolving the interval between stimulus onset and response (or display offset when no responses were required) with the canonical hemodynamic response function. Run means and movement parameters were included as covariates of no interest. The resulting beta-estimates were used to construct contrasts between the two conditions of each task, or for working memory, the contrast of rest against task as implicit baseline.

### Whole-brain univariate analysis

The between-condition contrasts that were used to examine DMN activity were: (1) belief > non-belief in the theory of mind task; (2) moral-personal dilemmas > non-moral dilemmas in the moral dilemmas task; (3) autobiographical memory > general knowledge in the autobiographical memory task; (4) landmarks > digits in the spatial imagery task; (5) self > other in the self/other adjective judgement task; and (6) rest > working memory.

A second level whole-brain analysis (one-sample t-test across subjects) was conducted on each of the six within-subject contrasts above, to obtain group activation maps for each contrast separately. Activation maps were thresholded at p < 0.05, controlling the false discovery rate (FDR; Benjamini and Yekutieli, 2001). A whole-brain analysis was conducted to examine individual participant activations for each of the six contrasts. For each voxel, we computed the number of participants with significant activation, applying FDR correction across all voxels of all participants (Heller et al., 2007). This resulted in a whole-brain map showing the number of participants with significant activation within each voxel. Based on the six random-effects analyses above, a similar map was constructed to show the number of significant task contrasts at each voxel (Heller et al., 2007). MRIcroN (Rorden et al., 2007) was used for visualization of whole-brain maps.

### Regions of interest and ROI analysis

A DMN mask was constructed using the 17 network parcellation from Yeo et al. (2011), concatenating networks 10, 15, 16, and 17. Networks 15, 16, and 17 largely corresponded to the three DMN subnetworks described in Andrews-Hanna (2012), which are the MTL subsystem, the dmPFC subsystem, and the core hubs. Network 10 was described in Yeo et al. (2011) as the orbital frontal-temporopolar network, which consists of temporopolar and orbital frontal regions. This network was added to the three DMN networks from Yeo et al. (2011) to include the vmPFC region described by Andrews-Hanna (2010). To create a single symmetrical volume, ROI masks (1 for voxels within the region; 0 outside) from the left and right hemispheres were combined using a logical OR operation, then projected back to both hemispheres. The combined network was then slightly smoothed (4 mm FWHM), and voxels with values > 0.5 after smoothing were retained. Finally, the combined network was parcellated into 20 smaller subregions by assigning each voxel to its closest DMN coordinate described by Andrews-Hanna et al. (2010). The coordinates are listed in Table 1. In cases where non-contiguous volumes were assigned to the same region, any volumes of < 45 voxels were discarded, and the remaining volume with center of mass closest to the Andrews-Hanna coordinate was chosen. The resulting ROIs are shown in Figure 2.

**Figure 2.**
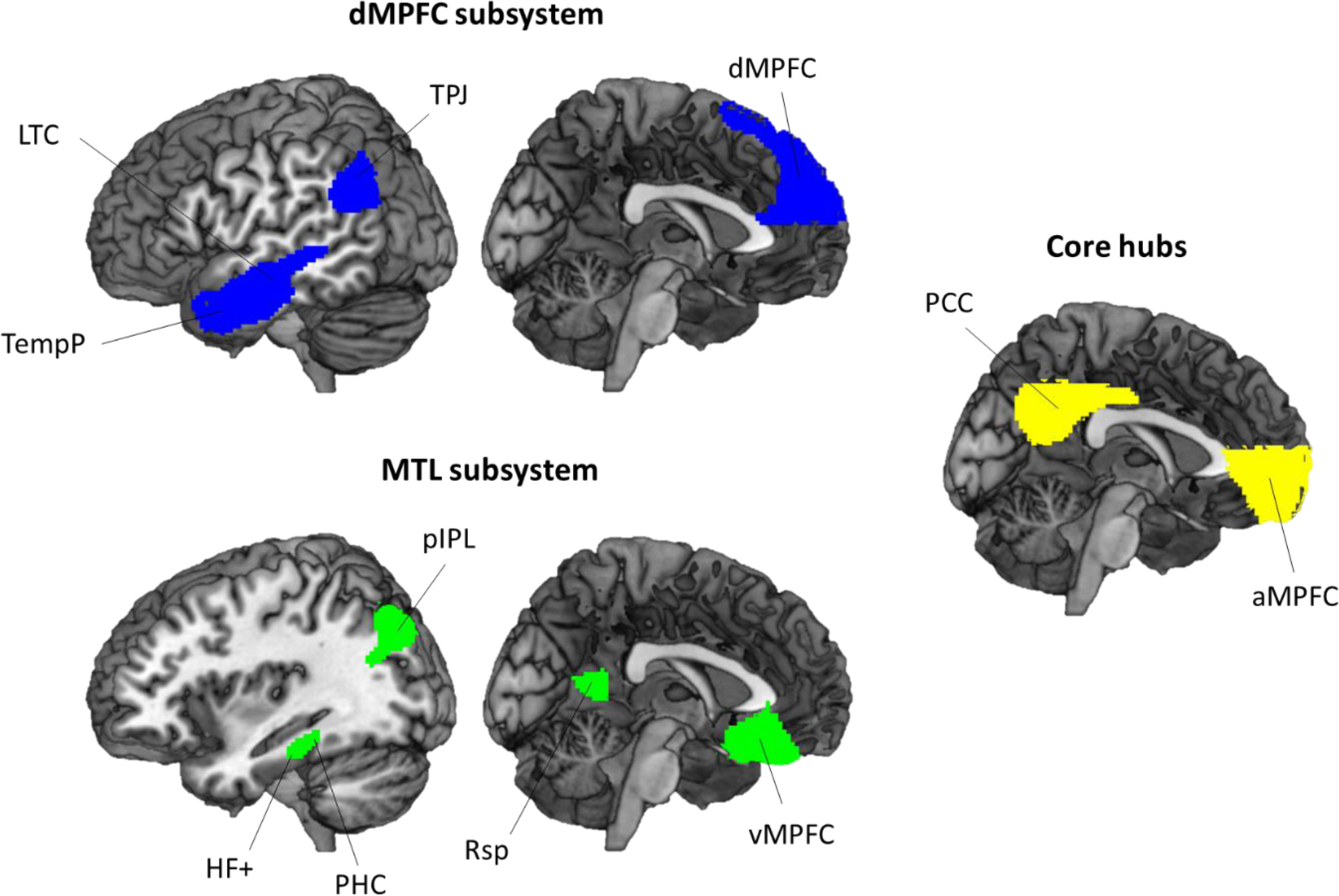
DMN ROIs used the in the current experiment. The ROIs are derived from networks 10, 15, 16, 17 described in the 17 network parcellation in Yeo et al. (2011) and devided according to coordinates described in Andrews-Hanna et al. (2010). Regions in blue are part of the dMPFC subsystem, and include the midline dMPFC and bilateral TPJ, LTC, and TempP. Regions in green are part of the MTL subsystem, and include the midline vMPFC and bilateral pIPL, Rsp, PHC, and HF+. The core hubs are represented in yellow, and include the bilateral aMPFC and PCC. For abbreviations see Table 1.

**Table 1.** Peak DMN coordinates described in Andrews-Hanna et al. (2010). Coordinates are based on the Montreal Neurological Institute coordinate system.

| Region | Abbreviation | x | y | z |
| --- | --- | --- | --- | --- |
| **dMPFC subsystem** |  |  |  |  |
| Dorsal medial prefrontal cortex | dMPFC | 0 | 52 | 26 |
| Temporal parietal junction | TPJ | -/+54 | -54 | 28 |
| Lateral temporal cortex | LTC | -/+60 | -24 | -18 |
| Temporal pole | TempP | -50/+50 | 14 | -40 |
| **MTL subsystem** |  |  |  |  |
| Ventral medial prefrontal cortex | vMPFC | 0 | 26 | -18 |
| Posterior inferior parietal lobule | pIPL | -/+44 | -74 | 32 |
| Retrosplenial cortex | Rsp | -/+14 | -52 | 8 |
| Parahippocampal cortex | PHC | -/+28 | -40 | -12 |
| Hippocampal formation | HF+ | -/+22 | -20 | -26 |
| **Core hubs** |  |  |  |  |
| Anterior medial prefrontal cortex | aMPFC | -/+6 | 52 | -2 |
| Posterior cingulate cortex | PCC | -/+8 | -56 | 26 |

For each task, the contrast between the two conditions was averaged within each ROI using the MarsBAR toolbox (Brett et al., 2002). For working memory, the relevant contrast was simply rest against implicit baseline (active task). Contrasts were tested against zero using two-tailed t-tests across subjects, corrected using FDR < 0.05 for multiple comparisons across ROIs. ROI × task ANOVAs were used to examine differences in ROI activity across different contrasts. Finally, the vector of contrast values from all tasks (six in total) was compared across ROIs. Distances between activation profiles for each pair of ROIs were calculated using 1 – Pearson’s r, and classical multidimensional scaling (MDS) was used to visualize the differences in activation pattern between ROIs as 2-dimensional distances.

### Task-wise multi-voxel pattern similarity

For each ROI, we wished to examine similarity of voxelwise activity patterns across the six tasks. For each participant, we extracted the beta-values for each contrast for each task, and compared the multivoxel patterns of these values between tasks. The similarity between each pair of tasks was measured by Pearson’s r, producing a symmetrical 6 × 6 matrix of similarities for each ROI. For each ROI, we quantified which regions showed (1) greater pattern similarity between the two tasks that required “introspection about mental states” (theory of mind and moral dilemmas), compared to similarity of these tasks to others, (2) greater pattern similarity between the two tasks that required “memory-based construction/simulation” (autobiographical memory and spatial imagery), compared to similarity of these tasks to others, (3) a relatively unique pattern for the self/other judgement task (greater similarity for task pairs not including self/other), and (4) a relatively unique pattern for rest (greater similarity for task pairs not including rest). To do this, we created four model similarity matrices based on these *a priori* groupings and evaluated fits to each ROI’s task similarity matrix using Kendall’s tau-a for each subject, as recommended when the model similarity matrix has ties (Nili et al., 2014). Correlations were tested against zero using 2-tailed t-tests across subjects, and all tests were corrected for multiple comparisons (FDR < 0.05) across the number of ROIs and models.

To compare patterns of task similarities between ROIs, we used vectors of between-task correlation from the above analysis (15 between-task correlations for each ROI). Similarly to the univariate analysis, distances between each pair of ROIs were calculated using 1 minus the correlation (Pearson’s r) between these vectors. Again, classical multidimensional scaling (MDS) was used for visualization.

## Results

### Behavioral results

Mean reaction times (RT) for all responses are summarized in Table 2. The first three subjects’ RTs for the working memory task were not recorded due to technical error and were excluded in the analysis. Mean accuracies for the theory of mind, mental imagery, and working memory tasks are also summarized Table 2, along with mean ratings of recollection and difficulty for the autobiographical memory task.

**Table 2.** Reaction times (RT), accuracies, and ratings of each condition (mean ± standard error).

|  | Theory of Mind |  | Moral Dilemmas |  | Autobiographical memory |  | Spatial imagery |  | Self/Other Adjective Judgement |  | Working memory |
| --- | --- | --- | --- | --- | --- | --- | --- | --- | --- | --- | --- |
|  | Belief | Non-belief | MPD | NMD | Memory | Knowledge | Landmarks | Digits | Self | Other | Working memory |
| RT (s) | 3.24 ± 0.03 | 3.10 ± 0.02 | 3.50 ± 0.03 | 3.26 ± 0.04 | 1.84 ± 0.02 | 1.90 ± 0.02 | 2.30 ± 0.02 | 3.02 ± 0.05 | 1.44 ± 0.01 | 1.47 ± 0.01 | 1.84 ± 0.02 |
| Accuracy (% correct) | 87.4 ± 6.2 | 91.1 ± 4.7 | N/A | N/A | N/A | N/A | 91.7 ± 5.7 | 84.3 ± 8.6 | N/A | N/A | 76.4 ± 5.0 |
| Rating | N/A | N/A | N/A | N/A | Recollection<br>3.65 ± 0.02<br>Difficulty<br>1.41 ± 0.02 | Recollection<br>1.36 ± 0.01<br>Difficulty<br>1.91 ± 0.02 | N/A | N/A | N/A | N/A | N/A |

Paired t-tests were conducted between the two conditions of the first five tasks, with no correction for multiple comparisons, to examine how well-matched each of the two conditions were within a task. There were no differences in reaction time between the pairs of conditions in the theory of mind, moral dilemmas, autobiographical memory, and self/other adjective judgement task (all |t|s < 1.45, all ps >= 0.16). In the spatial imagery task, RTs were shorter for the landmarks condition than for the digits condition (t = −2.74, p = 0.01). There were no differences in accuracy between the pairs of conditions in the theory of mind and spatial imagery task (both |t|s < 1.62, both ps >= 0.12). As expected, ratings of recollection were significantly greater in the autobiographical memory condition than in the general knowledge condition (t = 21.01, p < 0.001); autobiographical memory was also rated less difficult than general knowledge (t = −4.47, p = 0.001).

### Whole-brain univariate analysis

A whole-brain random effects analysis was conducted separately for each of the six contrasts of interest (Figure 3A; belief > non-belief; moral-personal dilemmas > non-moral dilemmas; autobiographical memory > general knowledge; landmarks > digits; self > other; and rest > task). Consistent with previous findings, the group analysis revealed many regions that are commonly associated with the DMN. In most tasks, we see activation in the medial prefrontal cortex (MPFC) and posterior medial cortex including PCC, precuneus, and Rsp, as well as temporal and parietal regions on the lateral surface, including pIPL, TPJ, and LTC. Activity for the self/other adjective judgement task was less typical of the DMN, though strongly activated a large portion of the MPFC.

**Figure 3.**
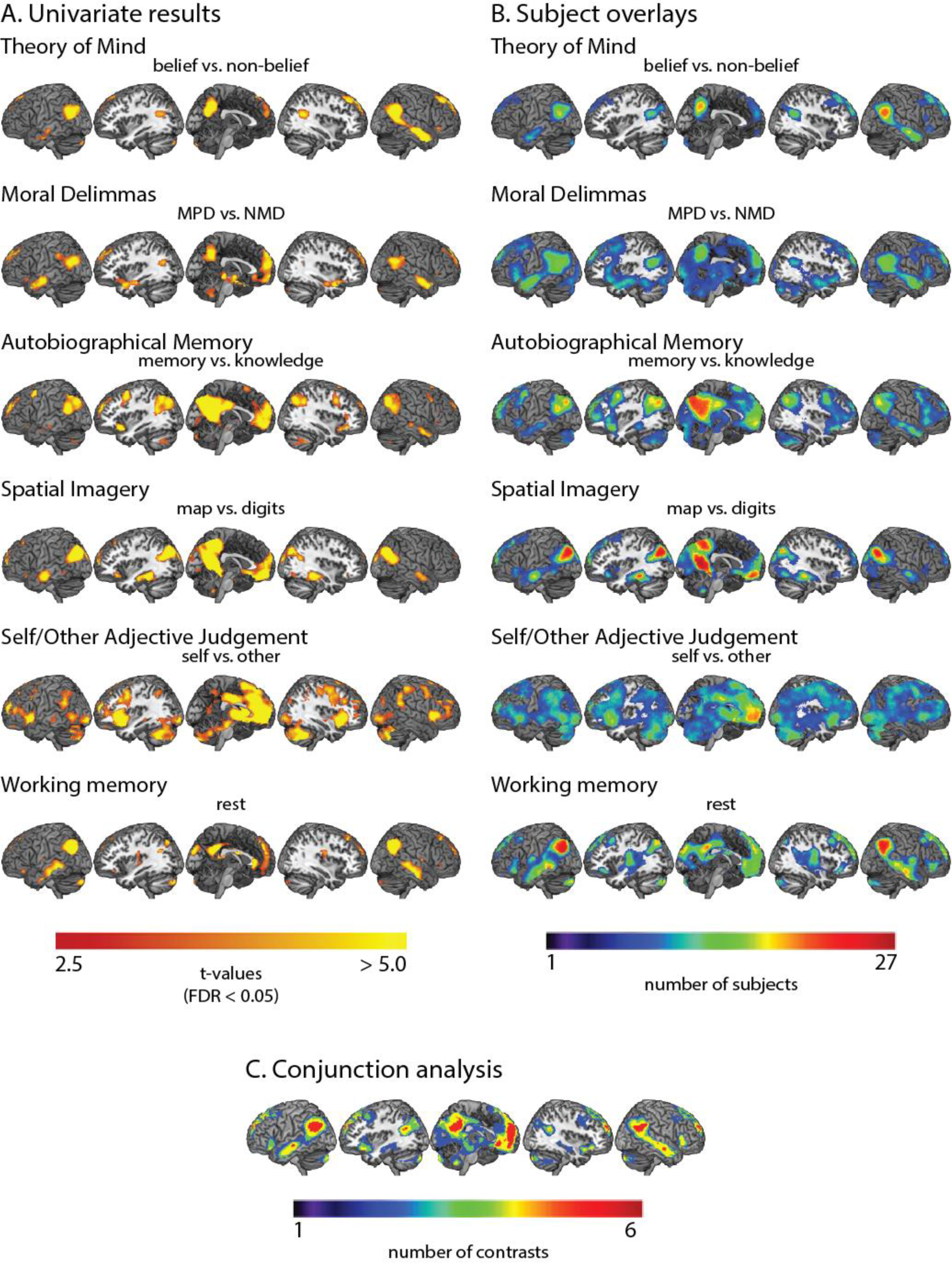
Univariate activity showing recruitment of the DMN network by all six tasks. (A) Whole-brain t-maps of the contrasts of interest in the six tasks. This includes belief > non-belief in the theory of mind task; moral-personal dilemmas > non-moral dilemmas in the moral dilemmas task; autobiographical memory > general knowledge in the autobiographical memory task; landmarks > digits in the spatial imagery task; self > other in the self/other adjective judgement task; and rest > task in the working memory task (working memory as implicit baseline). t-maps were thresholded at p < 0.05 (FDR corrected). (B) Overlay map of significant activations found in single subjects in the contrasts of interest. The color of each voxel represents the number of subjects that had significant activation in that voxel for a particular contrast, thresholded at 1 subject. (C) Overlay map of the number of significant contrasts from the six second-level analyses. The color of each voxel represents the number of contrasts that had significant activation in that voxel, thresholded at 2 contrasts.

To further quantify consistency across subjects, we computed a whole-brain overlay map for each task, where warmer colors indicate greater number of participants with significant activations (Figure 3B). The subject overlay map is largely consistent with the random effects results, as expected, but also indicates variability across participants.

Next, we identified regions that were consistently significantly activated across multiple contrasts (Figure 3C). No region was found to be active in all six contrasts after correcting for multiple comparisons (FDR < 0.05). However, several regions showed significant involvement in at least five contrasts. These include the MPFC (including dMPFC, aMPFC, and vMPFC), PCC, pIPL, TPJ, and parts of the LTC.

The results show that all six manipulations activated much of the DMN, and in particular, voxels within the MPFC, PCC, pIPL, TPJ, and LTC were significantly active for at least five manipulations. The theory of mind and moral dilemmas tasks showed strong activation of dMFPC, while the autobiographical memory and spatial imagery tasks showed peaks in vMPFC. These differences correspond to Andrews-Hanna’s (2012) observation of the dMPFC being involved in “introspection about mental states” and the vMPFC being involved in “memory-based construction/simulation”. Furthermore, the theory of mind and moral dilemmas tasks activated more anterior portions of the IPL than the autobiographical memory and spatial imagery tasks. This again corresponds to the separation of the TPJ (more anterior) and pIPL (more posterior) regions of the IPL, and matches their assignment to the dMPFC and MTL subsystems. The self > other contrast most consistently activated the MPFC across subjects, one of the core hubs identified by Andrews-Hanna (2012) to be responsive to “personally significant information”. However, the other hub region, the PCC, was only weakly activated. Our results show activity across much of the DMN for multiple contrasts, along with a degree of differentiation between dMPFC and MTL subsystems.

### ROI analysis of univariate activation level

For each of our six contrasts, profiles of activity across DMN ROIs are shown in Figure 4A(1). All contrasts were compared against zero using t-tests and were corrected for multiple comparisons with FDR < 0.05.

**Figure 4.**
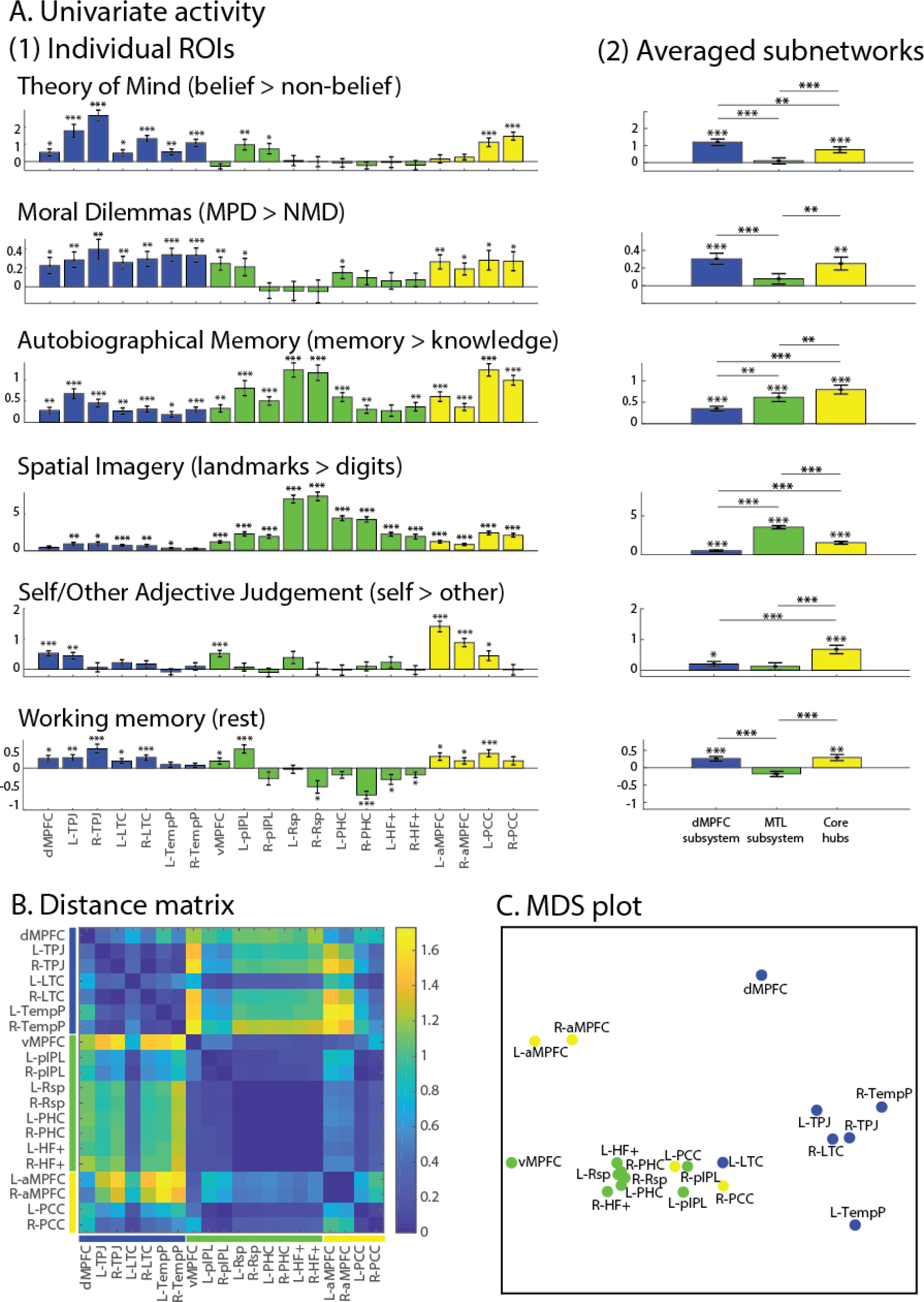
(A) DMN ROIs recruited by each condition within the 6 tasks. Error bars represent standard error. t-tests against zero were conducted for each contrast in each (1) ROI or (2) subnetwork. *** indicates p < 0.001, ** indicates p < 0.01, and * indicates p < 0.05 (all tests were corrected for multiple comparisons using FDR). (B) Dissimilarity matrix calculated using 1 – Pearson’s r between ROIs based on their activity profile across the 6 tasks. (C) Multidimensional scaling (MDS) to visualize the dissimilarity between regions.

Examined in detail, profiles suggest some of the anticipated differences between DMN regions, but also some surprises. As expected, theory of mind and moral dilemmas showed significant activation in most regions of the dMPFC and core networks. Activations were also seen in some regions of the MTL subsystem, however, including vMPFC, pIPL and PHC. Averaged contrasts within each network (Figure 4A(2)) showed significant activation just for the dMPFC subsystem and core. As anticipated, autobiographical memory and spatial imagery showed strong activations in the MTL subsystem, especially Rsp, and again in the core hubs, but significant activations were also seen in most dMPFC regions. Averaged within subsystems, the response of dMPFC was significantly lower than the other subsystems, but significantly greater than zero. For self-other, activations were more restricted, but included all three regions of the MPFC. Averaged within networks, this contrast was significant in the core and dMPFC subsystem, and, again as anticipated, strongest in the core subsystem. Unlike the previous four contrasts, core activation for self/other was stronger in aMPFC than in PCC. Perhaps surprisingly, the contrast of rest with working memory showed rather weak activations, significant only in the core and dMPFC subsystem, and significantly negative for some regions of the MTL network. Overall, these results provide broad support for the division into three subsystems, with the dMPFC subsystem especially involved in “introspection about mental states”, the MTL subsystem especially involved in “memory-based construction/simulation”, and the core hubs involved in all tasks but with particular sensitivity to “personally significant information”. At the same time, the results show that separation of networks is far from complete, with at least part of each network activated by every contrast. Within each network, there are also some clear variations in response. Notably, although the MTL subsystem as a whole was only active in the autobiographical memory and spatial imagery tasks, the pIPL and vMPFC were active for five of the six tasks, similar to the core hubs and dMPFC subsystem.

To compare profiles statistically, the data were entered into a repeated measures ROI (20) × task (6) ANOVA. Consistent with the different profiles suggested by Figure 4A(1), there was a strong interaction between ROI and task (F(95,2470) = 55.57, p < 0.001). There were also significant main effects for task (F(5,130) = 46.66, p < 0.001) and ROI (F(19,494) = 25.61, p < 0.001). The interaction in part reflects differences between the three subsystems, so we next repeated the ANOVA using the subsystem average profiles shown in Figure 4A(2). The significant interaction (F(10,260)=100.05, p < 0.001) confirms that this subnetwork grouping captures different functional profiles across the tasks. There were also main effects for networks (F(2,52) = 15.09, p < 0.001) and task (F(5,130) = 35.01, p < 0.001). We also wished to test for possible heterogeneity within each subsystem. To this end, ROI × task ANOVAs were repeated for each network separately. For the dMPFC subsystem, there was a significant interaction between ROI and task (F(30,780) = 9.21, p < 0.001), as well as main effects for ROI (F(6,156) = 27.54, p < 0.001) and task (F(5,130) = 12.50, p < 0.001). For the MTL subsystem, we also observed a significant interaction between ROI and task (F(40,1000) = 34.15, p < 0.001), as well as main effects for ROI (F(8,208) = 29.78, p < 0.001) and task (F(5,130) = 110.86, p < 0.001). Finally, there was also a significant interaction (F(15,390) = 22.92, p < 0.001) as well as main effects of ROI (F(3,78) = 25.42, p < 0.001) and task (F(5,130) = 12.87, p < 0.001) in the core hubs.

The distance matrix (Figure 4B), based on the dissimilarity of activation profiles for the 20 ROIs, showed distinct clusters. Profiles were largely similar for all regions in the dMPFC subsystem (Figure 4B, upper left), while dMPFC itself was somewhat separated from the cluster, being displaced towards aMPFC. In addition, the activation profile for L-LTC resembled the MTL as well as the other regions in the dMPFC subsystems. Regions in the MTL network also had largely similar profiles (Figure 4B, middle), but with other notable features. vMPFC resembled not only other MTL regions, but also aMPFC, while for pIPL, there was high similarity not only to other MTL regions, but also to much of the dMPFC subsystem and conspicuously also to PCC. Within the core regions, aMPFC had a relatively distinct profile, but was most similar to other frontal regions, while PCC instead showed results closely similar to those of pIPL, with similarity to all other regions except for aMPFC, dMPFC, and TempP.

These results are summarized in the MDS plot in Figure 4C. As expected, regions of the dMPFC network largely cluster together, but with dMPFC shifted towards other frontal regions. Regions of the MTL network are again close together, with vMPFC somewhat apart from the main cluster. PCC, instead of clustering with its partner core region, is placed between dMPFC and MTL networks, in a position close to pIPL. aMPFC occupies a position between the other two frontal regions, as perhaps expected from anatomical proximity.

### Task-wise multi-voxel pattern similarity

To compare the similarity of voxelwise activity patterns across tasks (e.g., belief > non-belief vs. self > other), we correlated patterns of beta-values across voxels, for each pair of tasks, within each ROI (Figure 5A). Four model similarity matrices were constructed to test (1) whether the two “introspection of mental states” tasks were especially similar, (2) whether the two “memory-based construction/simulation” tasks were especially similar, (3) whether the self/other adjective judgment task was especially dissimilar to other contrasts, and (4) whether rest > working memory was especially dissimilar to other contrasts. Results showed that the dMPFC subsystem (dMPFC, R-TPL, R-LTC, and R-TempP), as well as pIPL and PCC had strong pattern similarity between the two “introspection” tasks. On the other hand, the MTL subsystem (pIPL, Rsp, PHC, HF+), as well as aMPFC and PCC showed strong pattern similarity between the two “memory-based construction” tasks. Across many ROIs of the three subsystems, there was a strong tendency for the self > other pattern to be distinct from others (greater similarity for contrast pairs not involving self/other). Few regions, however, showed the rest > working memory pattern to be distinct from the others (only R-Rsp). Together, these data complement the findings in Figure 4. Though regions in each subsystem contain voxels responding to each contrast, the pattern of these activations is organized along the lines proposed by Andrews-Hanna (2012), with more dissimilar activation patterns for contrasts predominantly associated with different networks.

**Figure 5.**
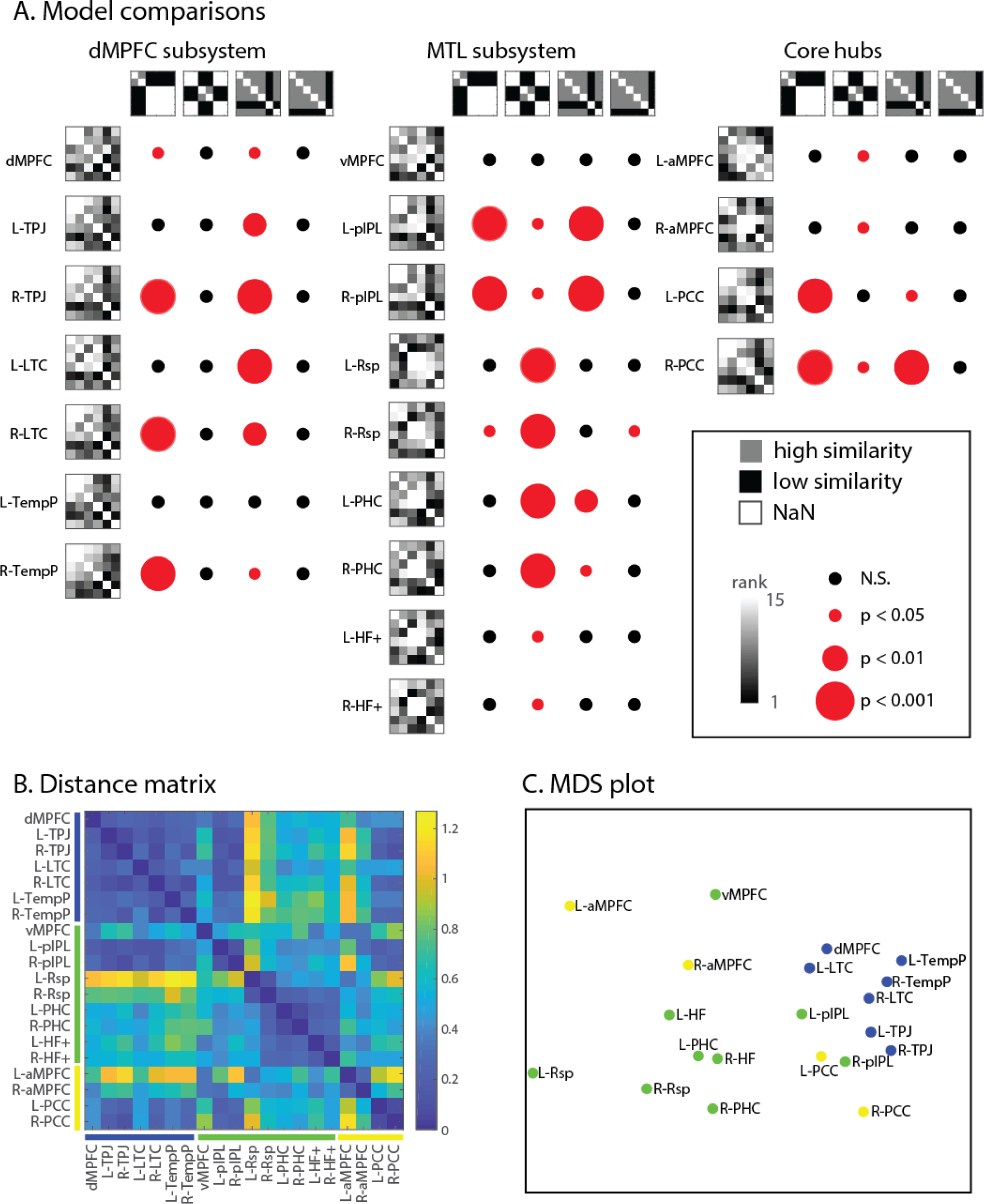
(A) Correlations between each pair of activation patterns in each ROI (subnetworks in columns). The upper row shows the four model similarity matrices (white indicates empty cells that were not used in the comparisons, grays indicate 1s, and black indicate 0s). In each matrix, tasks are ordered (top to bottom and left to right) as follows: theory of mind, moral dilemmas, autobiographical memory, spatial imagery, self/other and working memory. Leftmost columns show the average rank-transformed similarity matrices across subjects. Colored dots indicate the significance of Kendall’s tau-a correlation between each participant’s empirical and model similarity matrices tested against zero (corrected for multiple comparisons at FDR < 0.05). (B) Dissimilarity matrix calculated using 1 – Pearson’s correlation between ROIs based on their correlation profiles across 15 task pairs. (C) Multidimensional scaling (MDS) to visualize the dissimilarity between regions.

The distance matrix (Figure 5B) and MDS plot (Figure 5C), based on correlations of the pattern-similarity matrices shown in Figure 5A, showed distinct clusters, largely similar to those based on univariate activity profiles. The ROIs of the dMPFC subsystem clustered with each other, as did many of the MTL ROIs. Again, however, PCC and IPL regions clustered close together, between dMPFC and MTL clusters, and again, despite putative assignment to different networks, there was some similarity of the three MPFC regions.

## Discussion

Many complex cognitive processes have been linked to the DMN, supporting its role in high-level thought (Buckner and Carroll, 2007; Buckner et al., 2008; Spreng et al., 2009; Andrews-Hanna, 2012; Andrews-Hanna et al., 2014b). Among the most established of these cognitive functions are social, semantic, episodic, and self-relevant processing (Frith and Frith, 2006; Binder et al., 2009; McDermott et al., 2009; Spreng et al., 2009; Humphreys and Lambon Ralph, 2017). Recent findings suggest that the DMN consists of anatomically and functionally heterogeneous subsystems (Andrews-Hanna, 2012; Andrews-Hanna et al, 2014b; Yeo et al., 2011; Braga et al., 2017; Axelrod et al., 2017). Here, we used six diverse tasks to examine functional similarities and differences between DMN regions.

In many respects, our results matched the tripartite division proposed by Andrews-Hanna et al. (2010, 2014b; 2012). In terms of univariate activity, regions of the dMPFC subsystem had largely similar activity profiles (Figure 4B), with strong response to our two social tasks, consistent with a particular role in social cognition or introspection about mental states. A partial exception was dMPFC itself, whose activity profile was shifted towards that of aMPFC (Figure 4B, C). In addition to their strong response to social contrasts, however, dMPFC regions also showed some response to most other contrasts (Figure 4A). Thus, specialization was quantitative rather than qualitative. Analysis of multivoxel activity patterns also largely supported the proposals of Andrews-Hanna et al. (2010, 2014b; 2012), with regions of the dMPFC subsystem showing similar voxelwise activity patterns for our two social contrasts (Figure 5A), and again, largely similar profiles of between-task distances (Figure 5B, C).

Our results also support the proposal of an MTL subsystem, though with some caveats. In terms of univariate activity, regions of the MTL subsystem had largely similar activity profiles (Figure 4B), with strong response to the autobiographical memory and spatial imagery tasks, and in most cases little response to other contrasts (Figure 4A). The most conspicuous exceptions were vMPFC, whose activity profile was shifted towards that of aMPFC, and pIPL, which responded to most contrasts (Figure 4B, C). Analysis of multivoxel patterns showed a largely similar picture. For MTL regions except vMPFC, voxelwise activity patterns were especially similar for the memory and imagery contrasts (Figure 5A), and across all task pairs, there were largely similar profiles of between-task distances (Figure 5B, C). Again, though, the distance profile of pIPL was rather different, with some similarity to other regions of both MTL and dMPFC subsystems, and again, vMPFC was shifted towards aMPFC (Figure 5B, C).

Our results give less support to the concept of a midline core consisting of aMPFC and PCC. In terms of both univariate and multivariate activity, aMPFC was more similar to the adjacent dMPFC and vMPFC regions than to PCC. In terms of univariate activity, its strongest response was to the self-other contrast (Figure 4A). In contrast, both univariate and multivariate analyses placed PCC between dMPFC and MTL subsystems, with results closely similar to those of pIPL (Figure 4C, 4C). If anything, these results suggest pIPL and PCC as a DMN functional “core” (consistent with Buckner et al. (2008, 2009)), while MPFC regions show some dorsal-ventral gradient but also resemblances to one another, and relatively distinct profiles compared to the other ROIs, including PCC.

Some important caveats should be considered. Undoubtedly, our *a priori* ROIs would not match the exact functional regions of individual participants, meaning that results for adjacent regions will to some extent blur together. One region where this consideration could be especially significant is the inferior parietal lobule, represented here by pIPL and TPJ ROIs (Figure 2). Our univariate data agreed with the proposals of Andrews-Hanna et al. (2010b) in broadly separating pIPL and TPJ. At a finer scale, however, it is possible that pIPL should be further subdivided, as suggested by some functional connectivity data (Yeo et al., 2011). Blurring of functionally separate regions within the pIPL might contribute to our findings of similarity to both dMPFC and MTL subsystems, resembling PCC. Similar considerations apply to our finding of broad similarities between the three MPFC regions. Of particular relevance here are the results of Braga & Buckner (2017), who scanned four individuals 24 times using fMRI. The authors found that two distinct networks, showing resemblance with the dMPFC and MTL subsystems in Andrews-Hanna et al. (2010), could be identified in each individual. However, a unique finding from this study was that spatially juxtaposed regions of the two networks were found in each of the three MPFC regions: dMPFC, aMPFC, and vMPFC, which may be blurred together by spatial averaging in a group analysis. Despite these concerns, our results confirmed a dorsal-ventral gradient within the MPFC, with the dMPFC being more involved in tasks requiring “introspection of mental states” and vMPFC more involved in tasks requiring “memory-based construction/simulation”.

Other aspects of our results cannot be explained by spatial blurring. In particular, a conspicuous result was a significant response to non-social contrasts throughout most regions of the dMPFC subsystem, including those far from the MTL or core hubs. Along with the broad similarity of whole-brain maps for each contrast (Figure 3), apart from self > other, such results confirm partial, but not complete separation of response patterns for different DMN subsystems.

As noted earlier, several authors have proposed that the DMN represents broad features of a cognitive episode, situation or context (Hassabis and Maguire, 2007; Ranganath and Ritchey, 2012; Manning et al., 2014). Our results suggest both partial functional separation but also integration within this context representation. Matching many other findings (Andrews-Hanna et al., 2014a; Axelrod et al., 2017), our results link regions of the dMPFC subsystem to social cognition, and regions of the MTL subsystem to spatial or scene representation. To represent a cognitive episode, it is plausible that social and spatial representations are often integrated, for example to indicate who is where in the represented episode. Such integration may be achieved through communication between dMPFC and MTL subsystems, perhaps especially mediated by the pIPL and PCC. The self is also likely to be a core part of any episode representation, perhaps especially dependent on MPFC. In this way, the DMN acts partly as an integrated whole, but binding together aspects of the episode representation that are predominantly contributed by separate subregions.

Two other regions are worthy of further consideration. The first is the inferior frontal gyrus (IFG), which was not part of our *a priori* ROIs. Our whole-brain results (Figure 3A) showed that although IFG activity was weak in second-level analyses for most tasks (with the exception of self > other), a substantial minority of individual participants showed reliable recruitment for most tasks (Figure 3B). In the semantic literature, it has been shown that the semantic network, including the IFG, consists of many regions overlapping with the DMN (Binder et al., 2009; Noonan et al., 2013). In a dataset of 1000 participants, Yeo et al. (2011) identified the IFG as part of the dMPFC subsystem (Andrews-Hanna et al., 2014b). Given these findings, future studies should consider further the relation between the IFG and the DMN.

The second region requiring further consideration is the hippocampus. The hippocampal peak (HF+) defined in Andrews-Hanna et al. (2010) is not located in the hippocampus proper, but lies between the PHC and perirhinal cortex (PRC) (Moore et al., 2014; Ritchey et al., 2015; https://neurovault.org/collections/3731/). The PHC has been linked to the “posterior medial system”, a network closely related to the DMN, while the PRC has been linked to the “anterior temporal system”, along with the temporal poles and orbitofrontal cortex (Ranganath and Ritchey, 2012). The role of the current HF+ ROI is therefore unclear as it may span functionally heterogeneous regions. Another question is whether the hippocampus is part of the DMN at all. Our results show a mixed picture, as only some contrasts activated parts of the hippocampus. Although the hippocampus has been associated with episodic memory and spatial navigation (Maguire et al., 1998; Addis et al., 2007; Rugg et al., 2012; Brown et al., 2016), it has been proposed to play a different role from other regions in the MTL subsystem. In particular, the hippocampus may integrate information across the anterior temporal and posterior medial systems (Ranganath and Ritchey, 2012).

Our findings provide a mixed answer to the question of functional specialization within the DMN. On the one hand, there is evidence of a largely integrated whole, with similar whole-brain activity maps for multiple contrasts, and some response to every contrast in each of the proposed subsystems, supporting classical accounts (e.g. Buckner and Carroll, 2007; Spreng et al., 2009). On the other hand, there is partial functional separation, in close accord with the proposals of separate dMPFC and MTL subsystems (Andrews-Hanna et al., 2010, 2014a; Andrews-Hanna, 2012), though with remaining uncertainties over the concept of a midline core. Integrating social, spatial, self-related, and other aspects of a cognitive situation or episode, the DMN may provide the broad context for current mental activity.

## Acknowledgements

This work was supported by funding from the Medical Research Council (United Kingdom), program SUAG/002/RG91365. TW was supported by the Medical Research Council studentship and the Percy Lander studentship from Downing College.

